# A potential prokaryotic and microsporidian pathobiome that may cause shrimp white feces syndrome (WFS)

**DOI:** 10.1101/2021.05.23.445355

**Authors:** Anuphap Prachumwat, Natthinee Munkongwongsiri, Wiraya Eamsaard, Kanokwan Lertsiri, Timothy W. Flegel, Grant D. Stentiford, Kallaya Sritunyalucksana

## Abstract

White feces syndrome (WFS) in shrimp cultivation ponds is characterized by the occurrence of shrimp with abnormal, white intestines (midguts) combined with large floating mats of white, shrimp fecal strings. The etiology for WFS is complex, similar to diarrhea in humans. EHP-WFS is a type of WFS characterized by massive quantities of spores from the microsporidian parasite *Enterocytozoon hepatopenaei* (EHP) together with mixed, unidentified bacteria in the shrimp hepatopancreas, midgut and fecal strings. However, WFS does not always develop in shrimp with severe EHP infections in controlled laboratory challenges. Further, in EHP-WFS outbreak ponds, some shrimp show white midguts (WG) while others in the same pond show grossly normal midguts (NG). We hypothesized that comparison of the microbial flora between WG and NG from the same EHP-WFS pond would reveal probable combinations of microbes significantly associated with EHP-WFS. To test this hypothesis, we selected a pond exhibiting a severe EHP-WFS outbreak in cultivated *Penaeus vannamei* and used a combination of microscopic and microbial profiling analyses to compare WG and NG samples. By histology, EHP plasmodia and spores were confirmed in the hepatopancreas (HP) and midgut of WG and NG shrimp, but pathological severity and spore quantity was higher in the WG shrimp. In addition, intestinal microbiomes in WG shrimp were less diverse and had higher abundance of bacteria from the genera *Vibrio* and *Propionigenium. Propionigenium* quantity in the HP of WG shrimp was significantly higher (*P* = 1.08e-5) than in NG shrimp (4,506 *vs*. 3 copies /100 ng DNA, respectively). These findings supported our hypothesis by revealing two candidate bacterial genera that should be tested in combination with EHP as a potential eukaryote-prokaryote pathobiome that causes EHP-WFS in *P. vannamei*.

**Highlights:**

- White feces syndrome (WFS) shrimp often harbor the microsporidian *Enterocytozoon hepatopenaei* (EHP)
- The hepatopancreas (HP) and midgut of EHP-WFS shrimp had more EHP copies and spores than EHP-non.-WFS shrimp
- *Vibrio* spp., *Propionigenium* sp. and EHP dominated in HP microbiomes of EHP-WFS shrimp
- *Propionigenium* copy numbers were uniquely high in the HP of EHP-WFS shrimp
- EHP-WFS shrimp also showed intestinal microbiomes of reduced diversity but more heterogeneity

## 1. Introduction

Shrimp cultivation ponds exhibiting white feces syndrome (WFS) are characterized by the occurrence of shrimp with abnormal, white intestines (midguts) combined with floating mats of white, shrimp fecal strings. The contents of the midguts and fecal strings vary among WFS outbreak ponds, but also frequently contain a mixed bacterial component. These two features indicate that WFS has a complex etiology similar to that outlined for other syndromic conditions in shrimp (Kooloth Valappil et al., 2021) and for animal and plant diseases more generally (Bass et al., 2019).

One type of WFS is characterized by the massive transformation and sloughing of microvilli from epithelial cells of tubules of the shrimp hepatopancreas (HP). These sloughed microvilli aggregate in the tubule lumens as vermiform bodies called aggregated, transformed microvilli (ATM) that superficially resemble gregarines (Sriurairatana et al., 2014). They accumulate in masses at both the HP center and the midgut and are excreted as white fecal strings that float because of high fat content. The causal mechanism for ATM formation is still unknown. This type of WFS is of relatively infrequent occurrence because ATM, although frequently produced, do not often accumulate in sufficient quantity to cause WFS. Even when they do, the WFS is not associated with severe mortality or other serious production problems (Sanguanrut et al., 2018). In our experience, white midguts may also be caused by heavy gregarine infections, severe *Vibrio* infections and hemocytic enteritis caused by ingested blue-green algae (Anjaini et al., 2018; Somboon et al., 2012), but they are not usually associated with accumulation of floating fecal mats. Thus, reports of WFS that are not accompanied by at least microscopic confirmation cannot be ascribed to any particular causative agent.

The type of WFS examined in this study is characterized by the presence in the shrimp midgut and in white fecal strings of sloughed hepatopancreatic cells, tissue debris and massive quantities of spores from the microsporidian parasite *Enterocytozoon hepatopenaei* (EHP) (Tourtip et al., 2009). Shrimp exhibit white to yellow-golden intestines, loose exoskeletons, reduced feeding and retarded growth, high size variation, reduced average daily growth, elevated feed conversion ratios and sometimes mortality. This type of WFS (here referred to as EHP-WFS) was first reported from Vietnam (Ha et al., 2010), but it was soon realized that EHP is not always associated with WFS and that shrimp can recover from WFS but remain infected with EHP (Flegel, 2012; Tangprasittipap et al., 2013).

EHP-WFS is currently being reported from China (Shen et al., 2019; Wang et al., 2020), Southeast Asia (Caro et al., 2020; Desrina et al., 2020; Flegel, 2012; Ha et al., 2010; Sajiri et al., 2021; Tang et al., 2016) and South Asia (Rajendran et al., 2016). It typically occurs after 40 days of culture. In Thailand, EHP-WFS occurrence has significantly increased across all aquaculture regions in recent years, and it economically threatens Thai shrimp production due to combined losses from retarded growth and sometimes mortality.

Histopathology in the HP of EHP-WFS shrimp can be distinguished from that of usual EHP infections by the massive, simultaneous production and release of spores by cell lysis, together with sloughing of whole cells containing spores and sometimes unidentified bacterial cells. Altogether, this results in a loss of integrity of the HP tubule epithelium and may be accompanied by some shrimp mortality. The spores, sloughed cells and debris from lysed cells accumulate in the midgut making it and the fecal strings white. This process does not normally occur even with severe EHP infections in the laboratory where cells with pre-spore plasmodia greatly outnumber cells that produce spores (Chaijarasphong et al., 2020; Flegel, 2012). In addition, the cells that do lyse to release spores normally do so in a dispersed manner over time that allows for cell renewal and leaves the HP structure more-or-less intact. This allows for long-term infections that result in no external signs of disease but may cause retarded growth.

Since WFS is not always associated with EHP infections and cannot be reproduced in the laboratory in controlled challenge tests, it is possible that EHP may be a component cause of EHP-WFS, with EHP a necessary but insufficient solo cause of WFS. We previously hypothesized (Chaijarasphong et al., 2020) that WFS might be induced in shrimp with severe EHP infections via some unknown causative signal that induced simultaneous production of spores by all or most of the EHP plasmodia. This opened questions regarding the environmental factors and/or pathobiomes that might lead to EHP-WFS. It has previously been reported that EHP spores in white shrimp midguts, in white fecal strings and in severely infected HP tissue are frequently accompanied by bacterial cells of varied morphology (Tangprasittipap et al., 2013; Thitamadee et al., 2016). Thus, it is possible that the missing component-cause(s) of EHP-WFS may be bacterial in nature.

To investigate the possibility that the cause of EHP-WFS is a pathobiome that includes a eukaryote and prokaryote bacteria, we took advantage of the fact that during EHP-WFS outbreaks some of the shrimp in the pond show white midguts (WG) while others show grossly normal midguts (NG). We hypothesized that comparison of the microbial flora between WG and NG shrimp from the same EHP-WFS pond would reveal probable combinations of microbes significantly associated with EHP-WFS. To test this hypothesis, we used a combination of histopathological analysis and high-throughput 16S rRNA amplicon sequencing analysis of bacterial microbiomes to compare the HP and guts of WG and NG shrimp. Our comparative analyses showed distinct characteristics that separated WG and NG shrimp and revealed a significant association between EHP-WFS and dominant bacterial taxa of the genera *Vibrio* and *Propionigenium*.

## 2. Material and methods

### 2.1 Shrimp sample collection

A *P. vannamei* shrimp cultivation pond exhibiting a WFS outbreak was chosen because of the high burden of EHP accompanied by some shrimp mortality. The pond was located in Chanthaburi Province, Thailand (see Table S1). It was completely polyethylene-lined and was at 27 days of culture with shrimp average weight 8.70 g and average daily growth of 0.32 g/day. Samples were collected on the 5^th^ day after the WFS outbreak began. Shrimp with white midguts (WG) and shrimp with grossly normal (digestive tracks) guts (NG) were arbitrarily selected and subjected to comparative histopathological, molecular and microbiome analyses. Altogether, 15 WG and 15 NG shrimp were collected, 10 each for microbiome and molecular analyses and 5 each for histopathological examination. Samples were collected under the approved protocol No. BT 07/2561 from BIOTEC Institutional Animal Care and Use Committee (IACUC).

### 2.2 Microbiome and molecular analyses

Each shrimp was dissected to remove the gastrointestinal tract for separate collection of the stomach, hepatopancreas and midgut (intestine) in 1.5 ml tubes containing 500 µl of lysis buffer (50 mM Tris pH 9, 0.1M EDTA pH 8, 50 mM NaCl, 2% SDS, 100 μg/ml proteinase K) for DNA extraction using a QIAamp® DNA Mini Kit (Qiagen). DNA samples were used for bacterial profiling with high-throughput 16S rRNA amplicon sequencing and quantitative polymerase chain reactions.

### 2.3 Histopathological analysis

Shrimp specimens were prepared for histological examination by standard methods (Bell, Lightner, 1988). Briefly they were fixed in Davidson’s AFA fixative for 18-24 hours before transfer to 70% ethanol before tissue processing, embedding in paraffin, sectioning (4 µm thick) and staining with hematoxylin and eosin (H&E). Slides were examined using a Leica DM 750 equipped with a Leica ICC50 W digital camera.

### 2.4 High-throughput 16S rRNA amplicon sequencing

DNA samples were sent for quality control, Illumina library preparation and sequencing at Macrogen, Inc. (South Korea). Amplicons from the V3-V4 variable region of bacterial 16S rRNA were obtained using forward (5’-CCTACGGGNGGCWGCAG-3’) and reverse (5’-GACTACHVGGGTATCTAATCC-3’) primers (Herlemann et al., 2011) and used for sequencing library preparation with a Herculase II Fusion DNA Polymerase Nextera XT Index Kit V2. Library concentration and size distribution were quantified with TapeStation D1000 before sequencing with the Illumina MiSeq platform using the 2×300 paired end format.

### 2.5 Analysis of microbiomes

The raw sequencing reads were trimmed to remove primer sequences by Cutadapt (https://cutadapt.readthedocs.io/) and later processed using QIIME2 (version 2019.7.0) (Bolyen et al., 2019) with dada2 denoise-paired (Callahan et al., 2016) with truncated lengths of 280 and 235 base pairs for forward and reverse reads, respectively, to produce a set of amplicon sequence variants (ASVs). Taxonomic classification of ASVs was performed with USearch against an RDP database (Edgar, 2010). ASVs were imported into R and filtered for ASVs found in ≥2 samples and of either ≥1% or ≥0.1% abundance for further analyses. The analyses of ≥1% abundance ASVs are presented in the main text, whereas those of the ≥0.1% abundance ASVs are given in the Supplementary Information. Filtered ASV sets were processed with either a compositional data (CoDa) analysis approach (Gloor et al., 2017) that examines the ratios between ASVs or a standard count data analysis. For the CoDa approach, zero count ASVs were replaced using the zCompositions R package (Palarea-Albaladejo, Martín-Fernández, 2015), transformed with the centered log ratio transform after which a singular value decomposition (SVD) was applied for principal-component analysis (PCA) plots; differential abundance tests for ASVs were performed with the ALDEx2 v1.6.0 Bioconductor package using significantly abundant ASVs of an expected effect size difference of ≥1 (Fernandes et al., 2014). For standard count data analysis, we used phyloseq (McMurdie, Holmes, 2013) and microbiome (http://microbiome.github.com/microbiome) packages for alpha diversity index calculation and non-metric multidimensional scaling (NMDS) with a Bray-Curtis dissimilarity distance and EdgeR (McCarthy et al., 2012) or DESeq2 (Love et al., 2014) packages for differential abundance tests (FDR < 0.05). Additional graphics were plotted with vegan (https://github.com/vegandevs/vegan/), ggplot2 (https://ggplot2.tidyverse.org) and ggpubr (https://rpkgs.datanovia.com/ggpubr/) packages.

### Molecular quantification with quantitative polymerase chain reactions

Quantitative polymerase chain reactions (qPCR) were used to quantify copy numbers of EHP and selected *Propionigenium* taxa per 100ng of total DNA extracted. Each qPCR reaction was performed in a total volume of 20 µL, consisting of 10 µL 2X KAPA SYBR FAST qPCR Master Mix (KAPA Biosystems, USA), 0.2 µM of forward primer, 0.4 µL Low ROX, 100 ng of template DNA and a volume of water to the final volume 20 µL. Primers for EHP were described in Jaroenlak et al. (2016) and in Kanitchinda et al. (2020), whereas those for *Propionigenium* taxa were designed in this study - PG16S-F (5’-TGGACAATGGACCAAAAGTCTG-3’) and PG16S-R (5’-TTCAGCGTCAGTATTCATCCAG-3’). DNA templates for standard curve construction were derived from purified target fragments with the same sets of corresponding primers in different estimated copy numbers of ten-fold dilutions from 10^8^ to 10^2^ copies/1uL. Amplifications for qPCR measurement were carried out using a 7500 Fast Real-time PCR System (Applied Biosystems, USA) with the following conditions: for EHP, 2 min at 94 °C, followed by 40 cycles of 30 s at 94 °C, 30 s at 64 °C, and 30 s at 72 °C; and for *Propionigenium* taxa, 3 min at 95 °C, followed by 40 cycles of 15 s at 95 °C, 30 s at 55 °C, and 30 s at 72 °C. No template control and DNA samples of HP, midguts and standard curves were obtained in triplicate reactions. Melting curve and standard curve analyses evaluated specificity of the reactions to obtain estimated copy numbers of samples with an automatic software-assigned baseline and a manually-set threshold at 0.145 using the ABI PRISM® 7500 Sequence Detection System software (v2.3).

## 3. Results

### 3.1 WFS pond clinical signs and histopathology of shrimp gastrointestinal tracts

WG shrimp had whitish gastrointestinal tracts including the stomach, HP and entire intestine. They also exhibited loose and soft shells. In contrast, NG shrimp appeared grossly normal (Fig. S1). Histopathological examination of WG and NG shrimp revealed both shared and different abnormalities. Shared abnormalities included atrophied cells and EHP spores within the HP tubule epithelial cells and in epithelial cells of the midgut region located within the HP (Figs. 1 and 2). Focal lesions comprising shrimp hemocytes encapsulating of aggregated EHP spores were also observed in both groups (Fig. 1A). Specific characteristics observed in WG shrimp were 1) a higher prevalence of EHP plasmodia and spores within the HP and midgut epithelial cells (Figs. 1 and 2), and 2) a higher burden of free EHP spores, sloughed HP cells and rod-shaped bacterial cells in the midgut lumen (Fig. 2A and inset).

**Figure 1.**
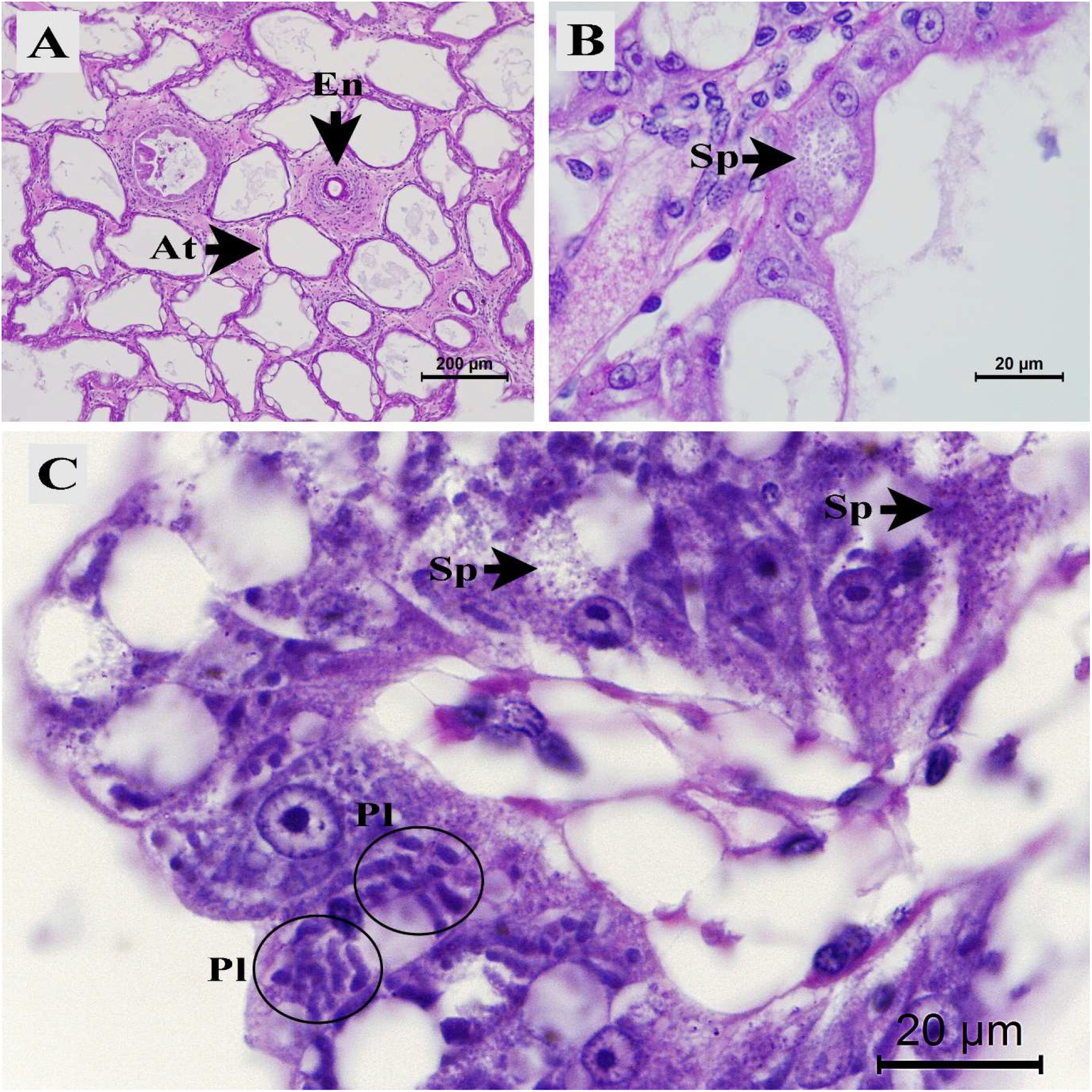
Photomicrographs of histological characteristics of white midgut (WG) and normal midgut (NG) shrimp hepatopancreatic tissues. Similar characteristics of WG and NG shrimp were **(A)** atrophied cells (At) of the hepatopancreatic epithelial tubule and hemocytic encapsulation (En), (**B**) EHP spores (Sp) in hepatopancreatic epithelial cells and (**C**) high prevalence of plasmodia (Pl) and spores (Sp) in hepatopancreatic tubule epithelia.

**Figure 2.**
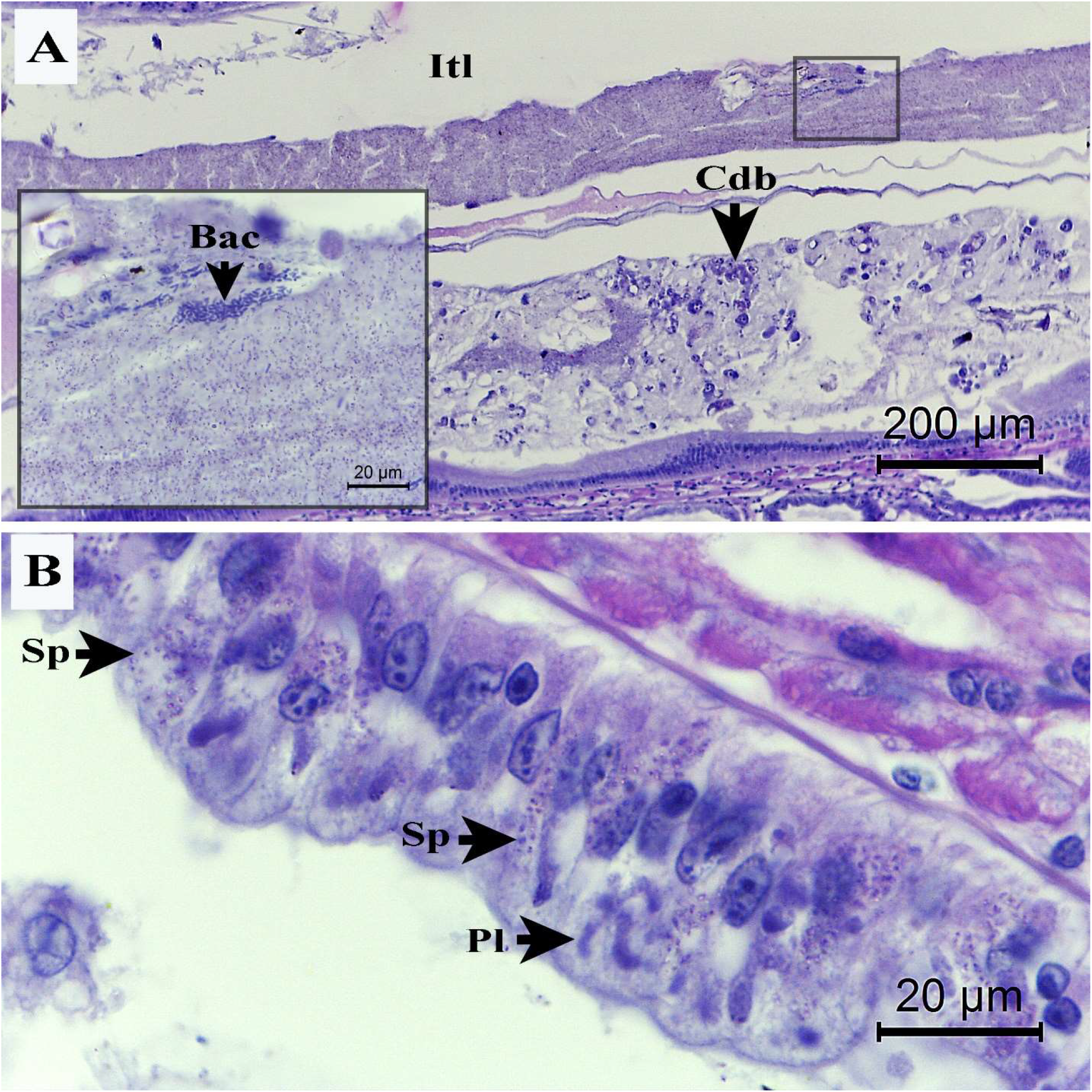
Photomicrographs of the midgut of white feces shrimp. (**A and inset**) Midgut lumen (Itl) containing HP epithelial cell debris (Cdb), colonies of rod-shaped bacteria (Bac) and masses of *Enterocytozoon hepatopenaei* (EHP) spores. (**B**) Spores (Sp) and plasmodia (Pl) of EHP-infected epithelial cells of the midgut. Note that the midgut epithelium is relatively normal and intact, despite the presence of EHP stages in some cells.

### 3.2 Comparison of intestinal microbiomes between WG and NG shrimp

Raw read pairs (5,974,443) were filtered to produce 2,254 amplicon sequence variants (ASVs) with QIIME2 DADA2 de-noise. These ASVs were filtered for only those found in ≥2 samples and of either ≥1% or ≥0.1% abundance. Initial examination of the two filtered datasets by principal-component analysis (PCA) and non-metric multi-dimensional scaling (NMDS) revealed that WG and NG samples had different bacterial profiles, except for one WG sample (F8) that closely clustered with NG samples (Supplementary Figs. S2, S3, S4 and S5; and Materials and Methods). PCA on the centered log-transformed data of the samples and the associated loadings for the ≥1% abundance ASV dataset (Fig. 3) revealed that intestinal bacterial communities between WG and NG shrimp differed markedly, except for the one WG shrimp sample (F8) that was more similar to the NG group. For the subsequent analyses, comparisons were made between the two groups: WG group of 12 sequenced library samples and NG group of 15 sequenced library samples (Supplementary Table S2), although similar trends were obtained when the F8 sample was included (data not shown).

**Figure 3.**
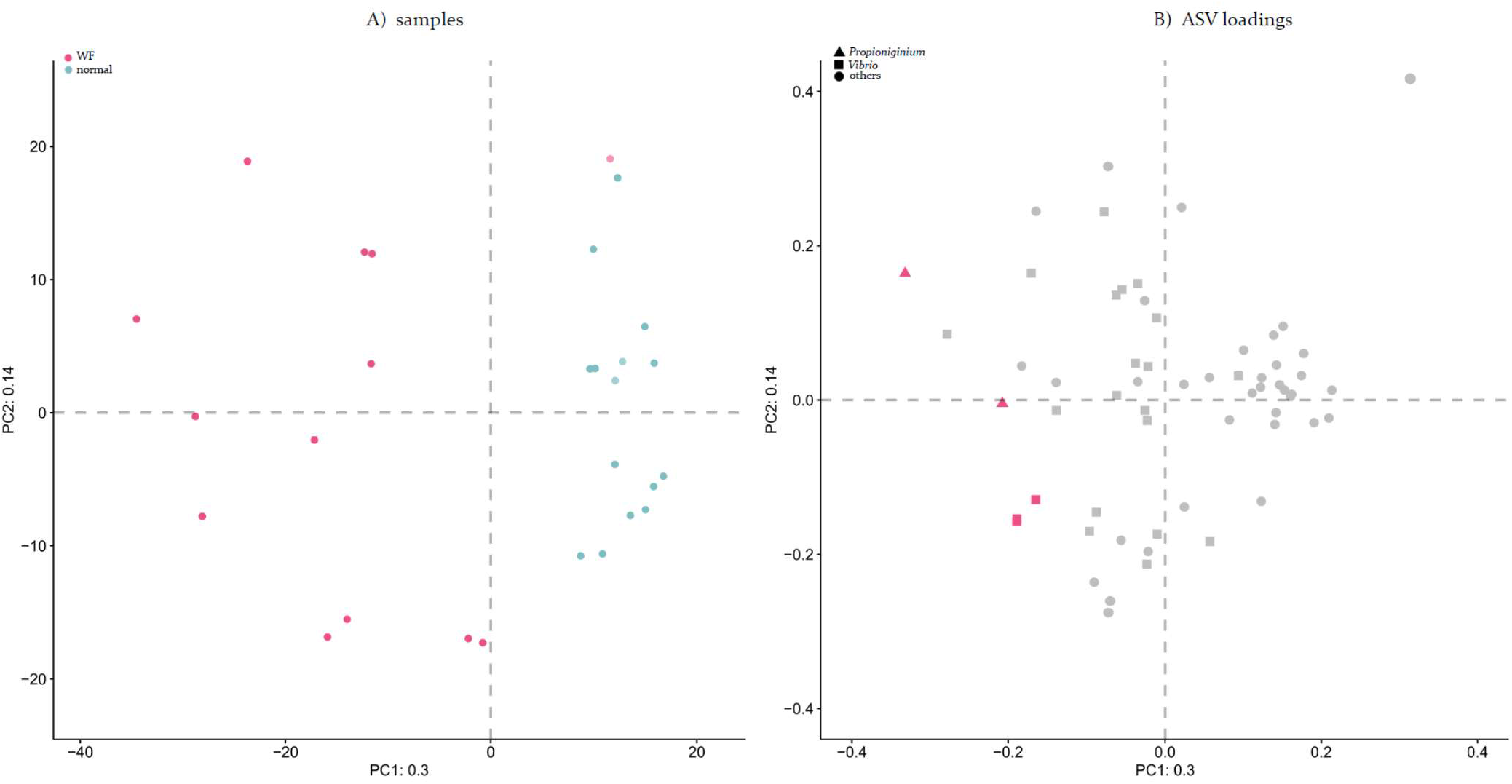
Compositional PCA plot of samples (A) and ASV loadings (B) for the ≥1% abundance ASV dataset (Materials and Methods). In panel **A**, each point is a sample [colored for WG (red) and NG (blue) shrimp groups; Table S1] and the distance between points is proportional to the multivariate difference between samples. Panel **B** shows the loadings for panel A in the same coordinate space, which represents the contributions of the ASVs to the separation of the samples. In this plot, each point is an ASV (shaped by taxonomic genus and colored by its assigned significantly higher abundance ASVs in the WG group (red) and the distance and direction from the origin to the point representing an ASV is proportional to the standard deviation of that ASV in the data set. The distance between one ASV and another is inversely proportional to their compositional association: points that are close together may have concordant relative abundances across all samples. The ability to directly interpret the plot is limited by the proportion of variance explained (30% on the first component and 14% on the second component).

A significantly lower alpha-diversity was observed in WG than in NG samples (Chao1’s Richness index, *P* = 2.5e-5; Shannon’s diversity index, *P* = 3.8e-7; Gini-Simpson index, *P* = 3e-3; Fig. 4). On average, distances among WG samples (on PCA and NMDS) were larger than those of the NG samples (Fig. 3 and Supplementary Figs. S2, S3, S4 and S5), suggesting more variation in bacterial communities between individual shrimp in the WG group than those in the NG group, i.e. the WG group was more heterogeneous.

**Figure 4.**
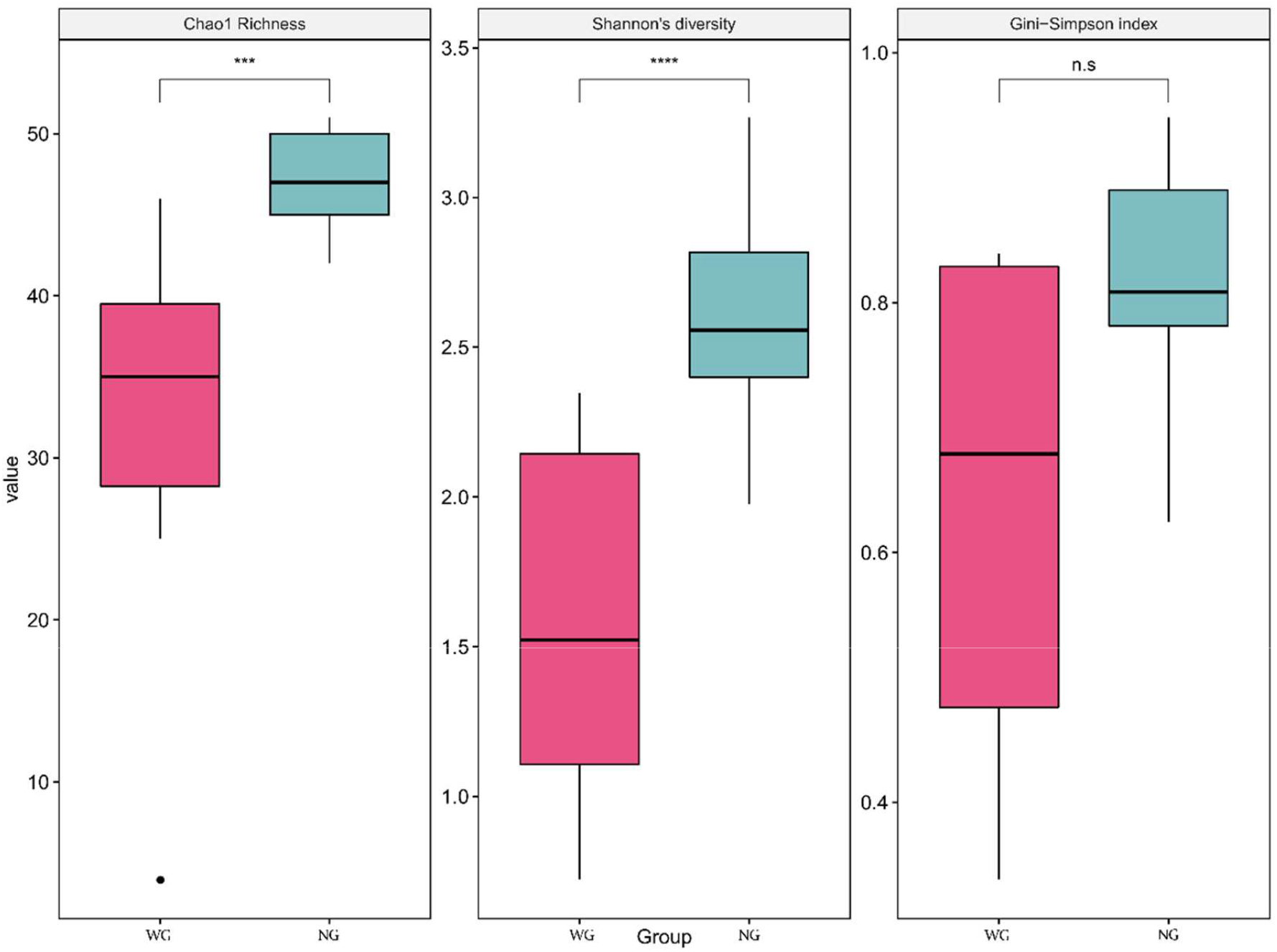
Comparison of alpha-diversity between the WG shrimp group (n = 12, red) and the NG shrimp group (n = 15, green) with Chao1’s Richness index, Shannon’s diversity index, and Gini-Simpson index. Significant differences are given by asterisks (n.s., P ≥ 0.05; ***, 0.0001 ≤ P < 0.001; ****, P < 0.0001).

To determine bacterial taxa associated with EHP-WFS shrimp, we analyzed for significantly over-represented ASVs in WG samples (see Materials and Methods). ASVs of the genera *Vibrio* and *Propionigenium* were found with significantly higher-fold changes across WG shrimp samples than NG samples (Figs. 3 and 5; see Materials and Methods). The average fold changes of over-represented abundances in WG over NG samples for genus *Vibrio* ASVs were 42.2 - 50.7 (3 ASVs), 5.2e7 (1 ASV), and 7.9e4 (1 ASV), from Aldex2, DESeq2 and EdgeR, respectively, while those for genus *Propionigenium* were 130 – 2004 (2 ASVs), 1.8e7 (1 ASV), 4.6e4 (1 ASV) from Aldex2, DESeq2 and EdgeR, respectively (Fig. 5). Similar ASVs relating to these *Vibrio* and *Propionigenium* taxa and some additional genera were also obtained with the other ASV datasets (see Materials and Methods and Supplementary Table S3).

**Figure 5.**
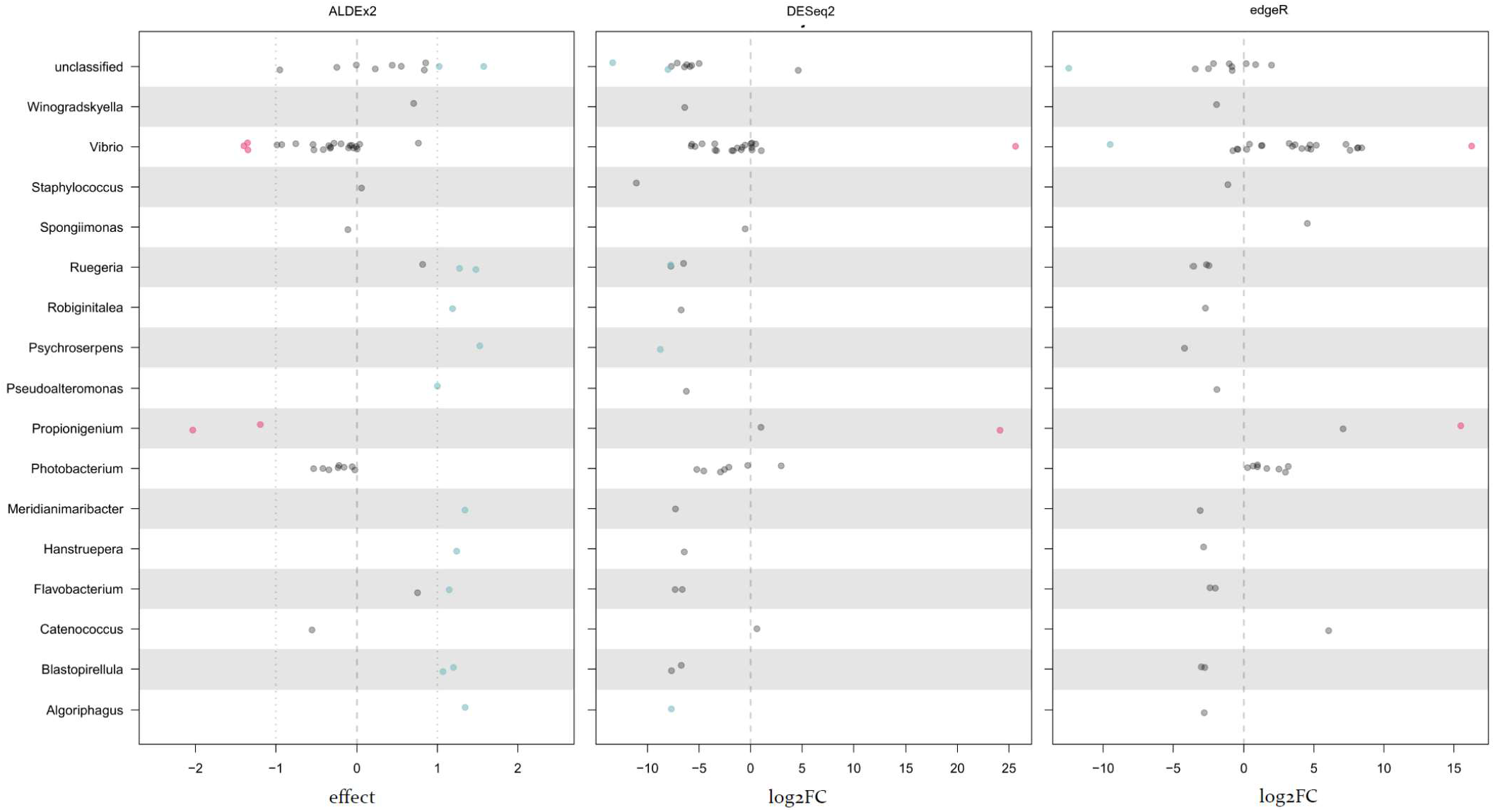
Differential relative abundance of ASVs binned by genus determined by (**A**) ALDEx2, **(B)** DESeq2 and (**C**) EdgeR. Points are colored as red or blue if they are significantly abundant in the WG or NG shrimp groups, respectively (Materials and Methods).

We focused on the significantly WG-over-represented ASVs of *Vibrio* and *Propionigenium* for further investigation relating to their significance in EHP-WFS. The sequences of significantly over-represented *Propionigenium* ASVs in WG samples all matched with *P. maris* with high identity (99.75% to those of *P. maris* or identical to those of uncultured and identified *Propionigenium* sp., which are likely strains of *P. maris*) such that specific primer sequences could be designed to compare abundance of *Propionigenium* in the shrimp gut and HP. However, significantly WG-over-represented *Vibrio* ASV sequences all matched records for multiple members of the *Vibrio harveyi* clade due to the short 16S rRNA region targeted. So also did the non-WG-over-represented *Vibrio* ASVs. Thus, it was not possible to make a species specific primer pair for comparative quantification of WG-over-represented *Vibrio*.

### 3.3 *High* Propionigenium *spp. levels in the hepatopancreas of WG but not NG shrimp*

To investigate different levels of EHP and *Propionigenium* between WG and NG shrimp, we carried out qPCR on the HP and midguts of all 10 WG and 10 NG shrimp. Copy number of both EHP and *Propionigenium* were significantly different between WG and NG shrimp in the HP but not in the midguts (Fig. 6 and Supplementary Table S2). Within the HP, there was a significantly higher copy number for *Propionigenium* in WG shrimp than in NG shrimp (medians of 4,506/100ng vs. 3/100ng DNA for WG and NG, respectively; *P* = 1.08e-5, Mann–Whitney U test; Fig. 6A). Specifically, all HP samples of NG shrimp had lower *Propionigenium* levels (< 50 copies) than the lowest standard copy number used (100 copies), implying that *Propionigenium* might be present at a low abundance or absent in the HP of NG shrimp. But all 10 HP samples from WG shrimp had > 100 copies of *Propionigenium*, suggesting its presence at high abundance in the HP of WG shrimp. For EHP within the HP, WG shrimp samples had a significantly higher copy numbers than did NG shrimp (medians of 37,573 and 24,966, respectively; *P* = 0.0432, Mann–Whitney U test; Fig. 6B), although both WG and NG shrimp had HP samples with similar maximum EHP levels (1.6e5 – 1.8e5 copies). Within midgut samples, both EHP and *Propionigenium* levels tended to be higher in WG than in NG (*Propionigenium* medians of 3,148 and 300 for WG and NG, respectively; *P* = 0.0524, Mann–Whitney U test; Fig. 6A; EHP medians of 13,928 and 6,900 for WG and NG, respectively; *P* = 0.4359, Mann–Whitney U test; Fig. 6B). Note that, contrary to its levels in the HP, *Propionigenium* levels in all midguts of NG shrimp were > 100 copies, where 100 copy number was the lowest standard copy number used in this experiment (Fig. 6A). When *Propionigenium* copy numbers were plotted against EHP copy numbers in shrimp HP samples (Fig. 6C), higher co-occurrence of *Propionigenium* and EHP was observed in WG shrimp while *Propionigenium* copies in the HP of NG shrimp were undetectable (< 50 copies/100ng DNA) by our qPCR assays (Fig. 6A and 6C).

**Figure 6.**
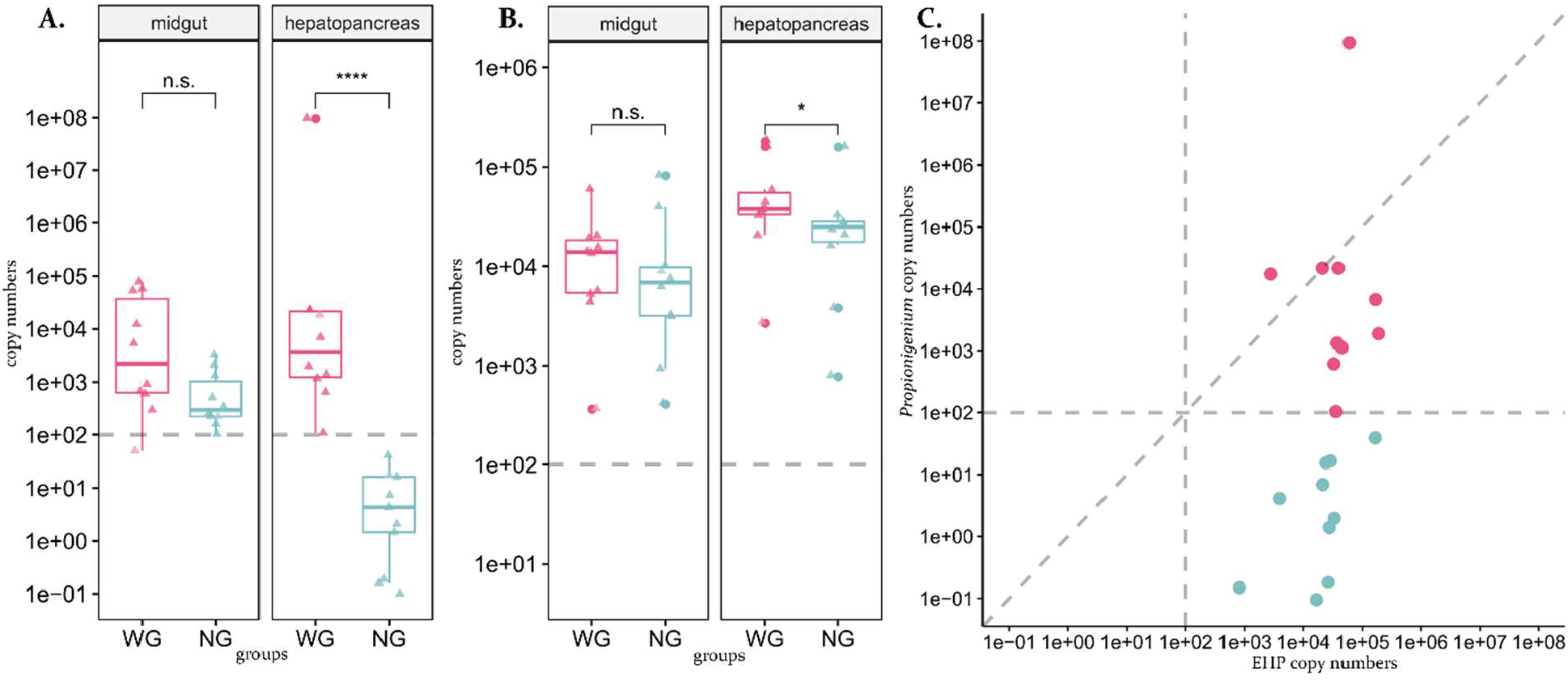
Copy numbers of *Propionigenium* (A) and EHP (B) in the 100 ng DNA samples of heapatopancreas and midguts between the WG and NG groups, and scatterplot (C) of the copy numbers of *Propionigenium* against those of EHP in the heapatopancreas samples. The points are colored red and blue for WG and NG groups, respectively. The estimated copy numbers were obtained by qPCR reactions described in the text.

## 4. Discussion

We have revealed the co-occurrence of EHP and distinctive bacterial communities which appear to contribute as a prokaryotic-eukaryotic pathobiome to cause the clinical manifestation of WFS in penaeid shrimp. Shrimp exhibiting these co-occurring microbial consortia in the HP displayed white gut (WG) characteristic of WFS, while others collected from the same shrimp pond but with normally colored guts (NG) lacked this pathobiome in the HP. With respect to gut histopathology, we confirmed earlier reports that WG shrimp exhibited more severe HP lesions characterized by higher numbers of spores and more tissue destruction (e.g., lysed, atrophied and sloughed cells) than did the NG shrimp (Figs. 1 and 2). We also proved that WG shrimp had significantly higher burdens of EHP than NG shrimp by qPCR counts. This was despite having similar maximum copy numbers for EHP by qPCR. In addition, high numbers of epithelial cells containing EHP plasmodia and/or spores were observed in the region of the midgut within the HP only in WG shrimp. Our examinations also confirmed earlier reports of rod-shaped bacterial cells being present together with the EHP spores.

In some of the WG and NG specimens, HP tissues showed lesions with hemocytic aggregation and encapsulation (Fig. 1A), but because bacteria were also present in many of the specimens, there was uncertainty as to whether these responses were induced by EHP or bacteria or both. Intracellular parasites usually do not elicit immune responses. For example, the microsporidian *Agmasoma penaei* in *P. monodon* muscle tissue rarely does (Flegel et al., 1992). Perhaps an inflammatory response can be initiated by tissue damage or cell lysis leading to release of intracellular parasite antigens. During microsporidiosis some insects such as lepidoptera and orthoptera display signs of cellular immunity by increased number of hemocytes, phagocytosis, encapsulation, nodule formation and melanization in infected tissues (Hoch et al., 2004; IaL et al., 2004; IuIa et al., 2000; Tokarev et al., 2007). However, some microsporidia species may escape or suppress host immunity for their advantage (Antúnez et al., 2009). Our histopathological examination also revealed a higher accumulation of rod-shape bacterial cells in the midgut lumen in WG than in NG shrimp, suggesting possible involvement of bacteria in conjunction with EHP in causing EHP-WFS.

Our high-throughput 16S rRNA amplicon sequencing analysis revealed that bacteria of the genera *Vibrio* and *Propionigenium* were significantly associated with WG shrimp (Figs. 3 and 5). It was subsequently confirmed that *Propionigenium* levels in HP and intestine samples of WG were higher than those in NG shrimp by qPCR (Fig. 6 and Supplementary Table S2). Similar comparisons could not be done with the dominant *Vibrio* species, the sequences of which were all related to the *Vibrio harveyi* clade (Darshanee Ruwandeepika et al., 2012; Ke et al., 2017; Urbanczyk et al., 2013). This was because the primers used for generic amplification of 16S rRNA yielded amplicons too short and too similarity to allow identification of individual *Vibrio* species within the clade. In this respect, we cannot discount the role of *Vibrio* taxa in the pathobiome of clinical EHP-WFS.. These associations between bacteria of the genera *Propionigenium* and *Vibrio* to EHP-WFS were observed and supported by both high-throughput 16S rRNA amplicon profiling and qPCR analyses.

Increased abundances of opportunistic *Vibrio* spp. measured by traditional plate counts have been reported in WFS ponds of both *P. monodon* and *P. vannamei* in many Asian countries. Specifically, reported *Vibrio* isolates from WFS shrimp gastrointestinal tracts and rearing water have been *V. harveyi, V. alginolyticus, V. parahaeolyticus, V. anguillarum, V. fluvialis, V. mimicus, V. vulnificus, V. damselae*, and *V. cholera* (Huang et al., 2020; Somboon et al., 2012; Supono et al., 2019; Wang et al., 2020). Some isolates have shown virulence in subsequent experimental bioassays by causing shrimp mortality but without WFS clinical signs (Wang et al., 2020). Using culture-independent approaches for high-throughput targeted amplicon or metagenomic shotgun sequencing, recent WFS studies have examined whether WFS intestinal microbial community assemblies differ from those of healthy shrimp. With WFS *P. vannamei* ponds in China and Indonesia, recent WFS microbiome studies reveals markedly different structures of WFS intestinal microbiomes that shift to gut “dysbiosis” with less diversity but more heterogenous bacterial composition than in healthy shrimp (Alfiansah et al., 2020; Hou et al., 2018; Huang et al., 2020; Wang et al., 2020). Shrimp gut dysbiosis has been observed in some EHP-WFS studies (Wang et al., 2020), but the other studies (Alfiansah et al., 2020; Hou et al., 2018; Huang et al., 2020) did not investigate EHP presence in their studied shrimp ponds. Importantly, key bacterial candidates associated with WFS were obtained by statistical analyses showing significantly more abundant bacterial taxa in WFS than in normal shrimp. These included taxa affiliated with *Vibrio, Candidatus* Bacilloplasma, *Aeromonas, Phascolarctobacterium, Ruminococcus, Rhodobacteraceae, Alteromonas, Marinomonas, Photobacterium, Pseudoalteromonas* and *Flavobacteraceae* (Alfiansah et al., 2020; Hou et al., 2018; Huang et al., 2020; Wang et al., 2020). Our microbiome analyses (Figs. 3 and 4) supported characteristics of lower bacterial diversities in WG samples and shifting of intestinal microbiome compositions to intestinal dysbiosis in WG shrimp. Our work added a bacterium from the genus *Propionigenium* to a list of WFS associated bacteria, specifically in EHP-WFS ponds exhibiting abnormal shrimp mortality.

The stark difference in absence of *Propionigenium* in the HP of NG shrimp but significant presence in the HP of WG shrimp in our study (Fig. 6A) was of particular interest. In contrast, its concentrations in the intestine of NG and WG were similar (Fig. 6A). It is possible that that progression of NG into WG shrimp might be associated to movement of *Propionigenium* (perhaps together with *Vibrio*) from the intestine to the HP.

The HP of healthy shrimp is usually devoid of bacteria, and presence of bacteria in the HP signifies poor health status (e.g., Vibriosis (Lightner, 1996)). The genus *Propionigenium* has not previously been associated with shrimp disease. The genus so far comprises two strictly anaerobic bacterial species (*P. maris* and *P. modestum*) that are capable of decarboxylating succinate to propionate for growth (Schink, 2006). They are found in marine habitats, typically in sediments. They are Gram negative, coccoid to ovoid or short rod-like cells with rounded ends (Schink, 2006). Of the two currently known species, our short 16S rRNA amplicon sequence showed the highest similarity to *P. maris*. Anoxic and metabolic conditions in shrimp intestines and HP might promote growth of *Propionigenium* during WFS progress.

Recently, succinic acid was one of metabolites found to be positively associated with WFS and with abundances of potential pathogenic bacteria such as *Vibrio*. Succinic acid is also a carbon source for *Propionigenium* that yields propionic acid. In addition, succinate supplemented feed in healthy shrimp can induce intestinal bacterial profile changes similar to those in WFS shrimp (Huang et al., 2020). Suggested avenues of further work include 1) tests on the possibility that propionic acid may induce spore formation and HP damage in *P. vannamei*, 2) work on the isolation and cultivation of *Propionigenium* from WFS shrimp for bioassays with EHP-infected shrimp and for species identification, and 3) epidemiological work to determine the risks factors (including the presence or absence of *Propionigenium* and *Vibrio* species) associated with WFS outbreaks.

## Acknowledgements

We would like to thank P. Wechprasit and W. Pattarayingsakul for their help in laboratory, D. Bass for helpful discussion on microbiome analysis, BIOTEC’s Biostatistics & Informatics Laboratory, K. Anekthanakul and Sai T. Y. A. for their programing and computational support. This work was financially supported by the Newton Institutional Links 2017, the Newton prize’s Chairman’s award, the International Collaborative Award (ICA\R1\180038) from the Royal Society (to GDS, Cefas/UK and KS, BIOTEC/Thailand). We also thank RDI management for National Strategic and Network Division (P19-51879), the National Science and Technology Development Agency (NSTDA).

## References

Alfiansah, Y.R., Peters, S., Harder, J., Hassenrück, C., Gärdes, A., 2020. Structure and co-occurrence patterns of bacterial communities associated with white faeces disease outbreaks in Pacific white-leg shrimp Penaeus vannamei aquaculture. Scientific reports. 10, 1–13.

Anjaini, J., Fadjar, M., Andayani, S., Agustin, I., Bayu, I., 2018. Histopathological in gills, hepatopancreas and gut of white shrimp (Litopenaeus vannamei) infected white feces disease (WFD). Research Journal of Life Science. 5, 183–194.

Antúnez, K., Martín‐Hernández, R., Prieto, L., Meana, A., Zunino, P., Higes, M., 2009. Immune suppression in the honey bee (Apis mellifera) following infection by Nosema ceranae (Microsporidia). Environmental microbiology. 11, 2284–2290.

Bass, D., Stentiford, G.D., Wang, H.-C., Koskella, B., Tyler, C.R., 2019. The pathobiome in animal and plant diseases. Trends in ecology & evolution. 34, 996–1008.

Bell, T.A., Lightner, D.V., 1988. A handbook of normal penaeid shrimp histology. World Aquaculture Society, Baton Rouge, Louisiana, USA.

Bolyen, E., Rideout, J.R., Dillon, M.R., Bokulich, N.A., Abnet, C.C., Al-Ghalith, G.A., Alexander, H., Alm, E.J., Arumugam, M., Asnicar, F., others, 2019. Reproducible, interactive, scalable and extensible microbiome data science using QIIME 2. Nature biotechnology. 37, 852–857.

Callahan, B.J., McMurdie, P.J., Rosen, M.J., Han, A.W., Johnson, A.J.A., Holmes, S.P., 2016. DADA2: high-resolution sample inference from Illumina amplicon data. Nature methods. 13, 581–583.

Caro, L.F.A., Mai, H.N., Pichardo, O., Cruz-Flores, R., Hanggono, B., Dhar, A.K., 2020. Evidences supporting Enterocytozoon hepatopenaei association with white feces syndrome in farmed Penaeus vannamei in Venezuela and Indonesia. Diseases of Aquatic Organisms. 141, 71–78.

Darshanee Ruwandeepika, H.A., Sanjeewa Prasad Jayaweera, T., Paban Bhowmick, P., Karunasagar, I., Bossier, P., Defoirdt, T., 2012. Pathogenesis, virulence factors and virulence regulation of vibrios belonging to the Harveyi clade. Reviews in Aquaculture. 4, 59–74.

Desrina, D., Prayitno, S.B., Haditomo, A.H.C., Latritiani, R., Sarjito, S., 2020. Detection of Enterocytozoon hepatopenaei (EHP) DNA in the polychaetes from shrimp ponds suffering white feces syndrome outbreaks. Biodiversitas Journal of Biological Diversity. 21.

Edgar, R.C., 2010. Search and clustering orders of magnitude faster than BLAST. Bioinformatics. 26, 2460–2461.

Fernandes, A.D., Reid, J.N., Macklaim, J.M., McMurrough, T.A., Edgell, D.R., Gloor, G.B., 2014. Unifying the analysis of high-throughput sequencing datasets: characterizing RNA-seq, 16S rRNA gene sequencing and selective growth experiments by compositional data analysis. Microbiome. 2, 1–13.

Flegel, T., Boonyaratpalin, S., Fegan, D., Guerin, M., Sriurairatana, S., 1992. High mortality of black tiger prawns from cotton shrimp disease in Thailand. Diseases in Asian Aquaculture I. 181, 197.

Flegel, T.W., 2012. Historic emergence, impact and current status of shrimp pathogens in Asia. Journal of invertebrate pathology. 110, 166–173.

Gloor, G.B., Macklaim, J.M., Pawlowsky-Glahn, V., Egozcue, J.J., 2017. Microbiome datasets are compositional: and this is not optional. Frontiers in microbiology. 8, 2224.

Ha, N., Ha, D., Thuy, N.T., Lien, V.T.K., 2010. Enterocytozoon hepatopenaei has been detected parasitizing tiger shrimp (Penaeus monodon) cultured in Vietnam and showing white feces syndrome. Agriculture and Rural Development: Science and Technology. 12, 45– 50.

Herlemann, D.P.R., Labrenz, M., Jurgens, K., Bertilsson, S., Waniek, J.J., Andersson, A.F., 2011. Transitions in bacterial communities along the 2000 km salinity gradient of the Baltic Sea. Isme J. 5, 1571–1579.

Hoch, G., Solter, L.F., Schopf, A., 2004. Hemolymph melanization and alterations in hemocyte numbers in Lymantria dispar larvae following infections with different entomopathogenic microsporidia. Entomologia experimentalis et applicata. 113, 77–86.

Hou, D., Huang, Z., Zeng, S., Liu, J., Wei, D., Deng, X., Weng, S., Yan, Q., He, J., 2018. Intestinal bacterial signatures of white feces syndrome in shrimp. Applied microbiology and biotechnology. 102, 3701–3709.

Huang, Z., Zeng, S., Xiong, J., Hou, D., Zhou, R., Xing, C., Wei, D., Deng, X., Yu, L., Wang, H., others, 2020. Microecological Koch’s postulates reveal that intestinal microbiota dysbiosis contributes to shrimp white feces syndrome. Microbiome. 8, 1–13.

IaL, V., IuS, T., IuIa, S., Glupov, V., 2004. Microsporidiosis in the wax moth Galleria mellonella (Lepidoptera: Pyralidae) caused by Vairimorpha ephestiae (Microsporidia: Burenellidae). Parazitologiia. 38, 239–250.

IuIa, S., IuS, T., IaL, L., Glupov, V., 2000. A morphofunctional analysis of the hemocytes in the cricket Gryllus bimaculatus (Orthoptera: Gryllidae) normally and in acute microsporidiosis due to Nosema grylli. Parazitologiia. 34, 408–419.

Jaroenlak, P., Sanguanrut, P., Williams, B.A.P., Stentiford, G.D., Flegel, T.W., Sritunyalucksana, K., Itsathitphaisarn, O., 2016. A Nested PCR Assay to Avoid False Positive Detection of the Microsporidian Enterocytozoon hepatopenaei (EHP) in Environmental Samples in Shrimp Farms. Plos One. 11.

Kanitchinda, S., Srisala, J., Suebsing, R., Prachumwat, A., Chaijarasphong, T., 2020. CRISPR-Cas fluorescent cleavage assay coupled with recombinase polymerase amplification for sensitive and specific detection of Enterocytozoon hepatopenaei. Biotechnol Rep (Amst). 27, e00485.

Ke, H.M., Prachumwat, A., Yu, C.P., Yang, Y.T., Promsri, S., Liu, K.F., Lo, C.F., Lu, M.J., Lai, M.C., Tsai, I.J., Li, W.H., 2017. Comparative genomics of Vibrio campbellii strains and core species of the Vibrio Harveyi clade. Sci Rep. 7, 41394.

Kooloth Valappil, R., Stentiford, G.D., Bass, D., 2021. The rise of the syndrome–sub‐optimal growth disorders in farmed shrimp. Reviews in Aquaculture.

Lightner, D.V., 1996. A handbook of shrimp pathology and diagnostic procedures for diseases of cultured penaeid shrimp.

Love, M.I., Huber, W., Anders, S., 2014. Moderated estimation of fold change and dispersion for RNA-seq data with DESeq2. Genome biology. 15, 1–21.

McCarthy, D.J., Chen, Y., Smyth, G.K., 2012. Differential expression analysis of multifactor RNA-Seq experiments with respect to biological variation. Nucleic acids research. 40, 4288–4297.

McMurdie, P.J., Holmes, S., 2013. phyloseq: an R package for reproducible interactive analysis and graphics of microbiome census data. PloS one. 8, e61217.

Palarea-Albaladejo, J., Martín-Fernández, J.A., 2015. zCompositions—R package for multivariate imputation of left-censored data under a compositional approach. Chemometrics and Intelligent Laboratory Systems. 143, 85–96.

Rajendran, K., Shivam, S., Praveena, P.E., Rajan, J.J.S., Kumar, T.S., Avunje, S., Jagadeesan, V., Babu, S.P., Pande, A., Krishnan, A.N., others, 2016. Emergence of Enterocytozoon hepatopenaei (EHP) in farmed Penaeus (Litopenaeus) vannamei in India. Aquaculture. 454, 272–280.

Sajiri, W.M.H.W., Borkhanuddin, M.H., Kua, B.-C., 2021. Occurrence of Enterocytozoon hepatopenaei (EHP) infection on Penaeus vannamei in one rearing cycle. Diseases of Aquatic Organisms. 144, 1–7.

Sanguanrut, P., Munkongwongsiri, N., Kongkumnerd, J., Thawonsuwan, J., Thitamadee, S., Boonyawiwat, V., Tanasomwang, V., Flegel, T.W., Sritunyalucksana, K., 2018. A cohort study of 196 Thai shrimp ponds reveals a complex etiology for early mortality syndrome (EMS). Aquaculture. 493, 26–36.

Schink, B., 2006. The genus Propionigenium. in: Dworkin, M.F.S.R.E.S.K.S.E. (Ed.), The Prokaryotes. Springer, New York, NY, pp. 3948-3951.

Shen, H., Qiao, Y., Wan, X., Jiang, G., Fan, X., Li, H., Shi, W., Wang, L., Zhen, X., 2019. Prevalence of shrimp microsporidian parasite Enterocytozoon hepatopenaei in Jiangsu Province, China. Aquaculture International. 27, 675–683.

Somboon, M., Purivirojkul, W., Limsuwan, C., Chuchird, N., 2012. Effect of Vibrio spp. in white feces infected shrimp in Chanthaburi, Thailand. Journal of Fisheries and Environment. 36, 7–15.

Sriurairatana, S., Boonyawiwat, V., Gangnonngiw, W., Laosutthipong, C., Hiranchan, J., Flegel, T.W., 2014. White feces syndrome of shrimp arises from transformation, sloughing and aggregation of hepatopancreatic microvilli into vermiform bodies superficially resembling gregarines. PloS one. 9, e99170.

Supono, S., Wardiyanto, W., Harpeni, E., 2019. Identification of Vibrio sp. as a cause of white feces diseases in white shrimp Penaeus vannamei and handling with herbal ingredients in East Lampung Regency, Indonesia. AACL Bioflux. 12, 417–425.

Tang, K.F., Han, J.E., Aranguren, L.F., White-Noble, B., Schmidt, M.M., Piamsomboon, P., Risdiana, E., Hanggono, B., 2016. Dense populations of the microsporidian Enterocytozoon hepatopenaei (EHP) in feces of Penaeus vannamei exhibiting white feces syndrome and pathways of their transmission to healthy shrimp. Journal of invertebrate pathology. 140, 1–7.

Tangprasittipap, A., Srisala, J., Chouwdee, S., Somboon, M., Chuchird, N., Limsuwan, C., Srisuvan, T., Flegel, T.W., Sritunyalucksana, K., 2013. The microsporidian Enterocytozoon hepatopenaei is not the cause of white feces syndrome in whiteleg shrimp Penaeus (Litopenaeus) vannamei. BMC veterinary research. 9, 1–10.

Thitamadee, S., Prachumwat, A., Srisala, J., Jaroenlak, P., Salachan, P.V., Sritunyalucksana, K., Flegel, T.W., Itsathitphaisarn, O., 2016. Review of current disease threats for cultivated penaeid shrimp in Asia. Aquaculture. 452, 69–87.

Tokarev, Y.S., Sokolova, Y.Y., Entzeroth, R., 2007. Microsporidia–insect host interactions: Teratoid sporogony at the sites of host tissue melanization. Journal of invertebrate pathology. 94, 70–73.

Tourtip, S., Wongtripop, S., Stentiford, G.D., Bateman, K.S., Sriurairatana, S., Chavadej, J., Sritunyalucksana, K., Withyachumnarnkul, B., 2009. Enterocytozoon hepatopenaei sp. nov.(Microsporida: Enterocytozoonidae), a parasite of the black tiger shrimp Penaeus monodon (Decapoda: Penaeidae): Fine structure and phylogenetic relationships. Journal of invertebrate pathology. 102, 21–29.

Urbanczyk, H., Ogura, Y., Hayashi, T., 2013. Taxonomic revision of Harveyi clade bacteria (family Vibrionaceae) based on analysis of whole genome sequences. Int J Syst Evol Microbiol. 63, 2742–2751.

Wang, H., Wan, X., Xie, G., Dong, X., Wang, X., Huang, J., 2020. Insights into the histopathology and microbiome of Pacific white shrimp, Penaeus vannamei, suffering from white feces syndrome. Aquaculture. 527, 735447.

